# A galling insect activates plant reproductive programs during gall development

**DOI:** 10.1101/383851

**Authors:** Jack C. Schultz, Patrick P. Edger, Mélanie J.A. Body, Heidi M. Appel

## Abstract

Some insects can redirect plant development to form unique organs called galls, which provide these insects with unique, enhanced food and protection from enemies and the elements. Many galls resemble flowers or fruits, suggesting that elements of reproductive development may be involved. We tested this hypothesis using RNA sequencing (RNAseq) to quantify the transcriptional responses of wild grapevine (*Vitis riparia* Michx.) leaves to a galling parasite, phylloxera *(Daktulosphaira vitifolia* (Fitch 1855)). If development of reproductive structures is part of gall formation, we expected to find significantly elevated expression of genes involved in flower and/or fruit development in developing galls as opposed to ungalled leaves. We found that reproductive gene ontology (GO) categories were significantly enriched in developing galls, and that expression of many candidate genes involved in floral development were significantly increased, particularly in later gall stages. The patterns of gene expression found in galls suggest that phylloxera exploits vascular cambium to provide meristematic tissue and redirects leaf development towards formation of carpels. The phylloxera leaf gall appears to be phenotypically and transcriptionally similar to the carpel, due to the parasite hijacking underlying genetic machinery in the host plant.

## Introduction

Plant galls are unique organs formed in response to a parasite, which may be a virus, fungus, bacterium, nematode, or arthropod^1^. The host’s development and resource allocation are modified to provide shelter and nutritionally superior food for the parasite^1,2^. Bacteria (e.g., *Agrobacterium*), elicit galls by integrating genes into the plant genome^1^, but their galls are best described as disorganized neoplasms. In contrast, arthropod galls are often as highly organized and complex as normal plant organs^3, 4, 5^. And arthropods do not genetically transform plant cells; removing the parasite halts gall growth and development, supporting the view that arthropods direct gall development chemically^6,7,8^, How this is done and the chemical cues involved remain poorly understood.

As Darwin^9^ noted, insect gall phenotypes often resemble plant reproductive organs. Both flower carpels and galls are structures that envelope, protect, and provide access to specialized food resources for an ‘alien’ organism: the plant embryo or insect. Like developing fruits, galls are usually strong sinks^10,11^ and provide nutritive cells rich in carbohydrates and proteins for the insect^3^ much as endosperm provides nutrition to plant embryos. A sclerotized capsule often protects the insect or plant embryo, and the surrounding cortex and epidermis can contain defensive chemistry^2,12^. These phenotypic traits develop under the direction of the insect or embryo using chemical signals (hormones, in the case of the embryo) and are encoded by a set of transcriptionally co-regulated genes^13^.

We examined the hypothesis that a galling insect, grape phylloxera *(Daktulosphaira vitifolia* (Fitch 1855)) co-opts portions of flower and/or fruit transcriptional pathways to produce a gall with the necessary attributes^14^. While some galls occur on flowers or fruits, most – including phylloxera - develop on vegetative tissues. Thus, assessing the degree to which a developing gall’s transcriptome diverges from that of the vegetative tissue on which it develops and specifically, the degree to which the gall’s transcriptome is reproductive, should indicate whether and to what degree the insect hijacks the underlying reproductive developmental programs in the host plant.

Grape phylloxera is a native North American aphid relative that elicits complex galls on abaxial leaf surfaces, and causes swelling on roots when feeding there. We employed RNA sequencing to characterize the transcriptome of this gall and the leaves on which it develops, sampling at four developmental intervals (Fig. 1.). We confirmed that the insect reprograms leaf cell transcriptomes to direct gall development^15^. We asked whether genes typical of floral organs, from the decision to flower through meristem establishment and floral organ formation^16^, were significantly enriched among genes differentially expressed in the gall compared to the leaf., Results confirmed that phylloxera gall development involves portions, but not all, of the floral developmental programs in grapevine.

**FIGURE 1.**
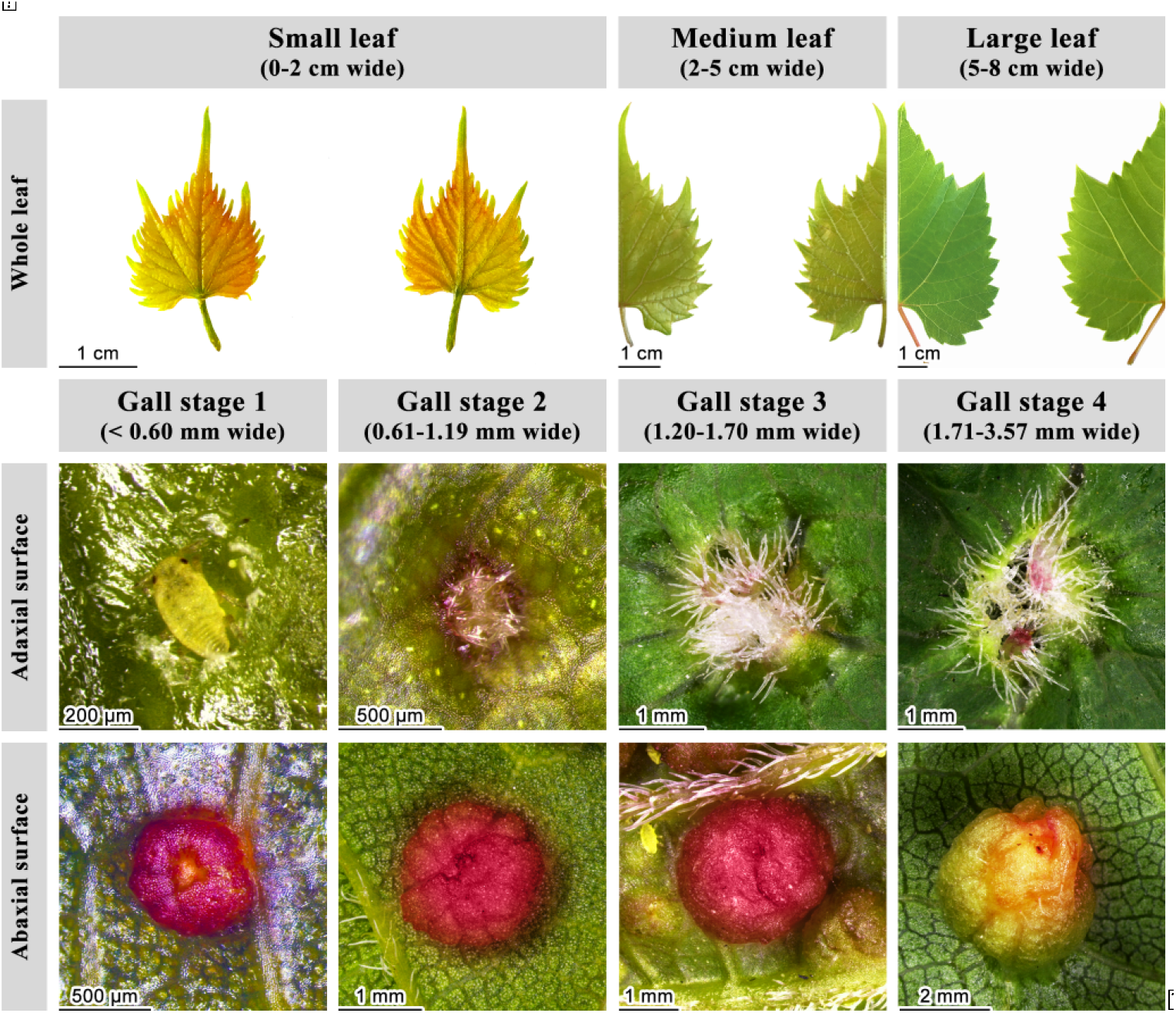
Gall stages sampled and the stage-matched leaves on which they occurred. The female is still visible at stage 1, but disappears as adaxial leaf tissue grows over her, while the sack-like gall expands beneath her. Very few galls are initiated on leaves wider than 1.2mm.

## Results

### Gall and ungalled leaf transcriptomes diverge significantly as the gall develops

We extracted RNA from phylloxera leaf galls at 4 intervals as they developed (Fig 1). Aligning reads to the *Vitis vinifera* genome (Version 12x; Phytozome Version 7, Joint Genome Institute) allowed us to identify 26,346 grape transcripts expressed in either gall or leaf or both. Of these 11,094 were differentially expressed (>1.5-fold, P<0.01) at least once in galls compared with ungalled leaves (Fig. 2.). Because the *Vitis* genome is not yet fully functionally annotated, we integrated *Vitis* transcripts with *Arabidopsis thaliana* TAIR v.9 functional annotations. This process produced 11,094 differentially-expressed transcripts we could potentially use for functional evaluation.

**FIGURE 2.**
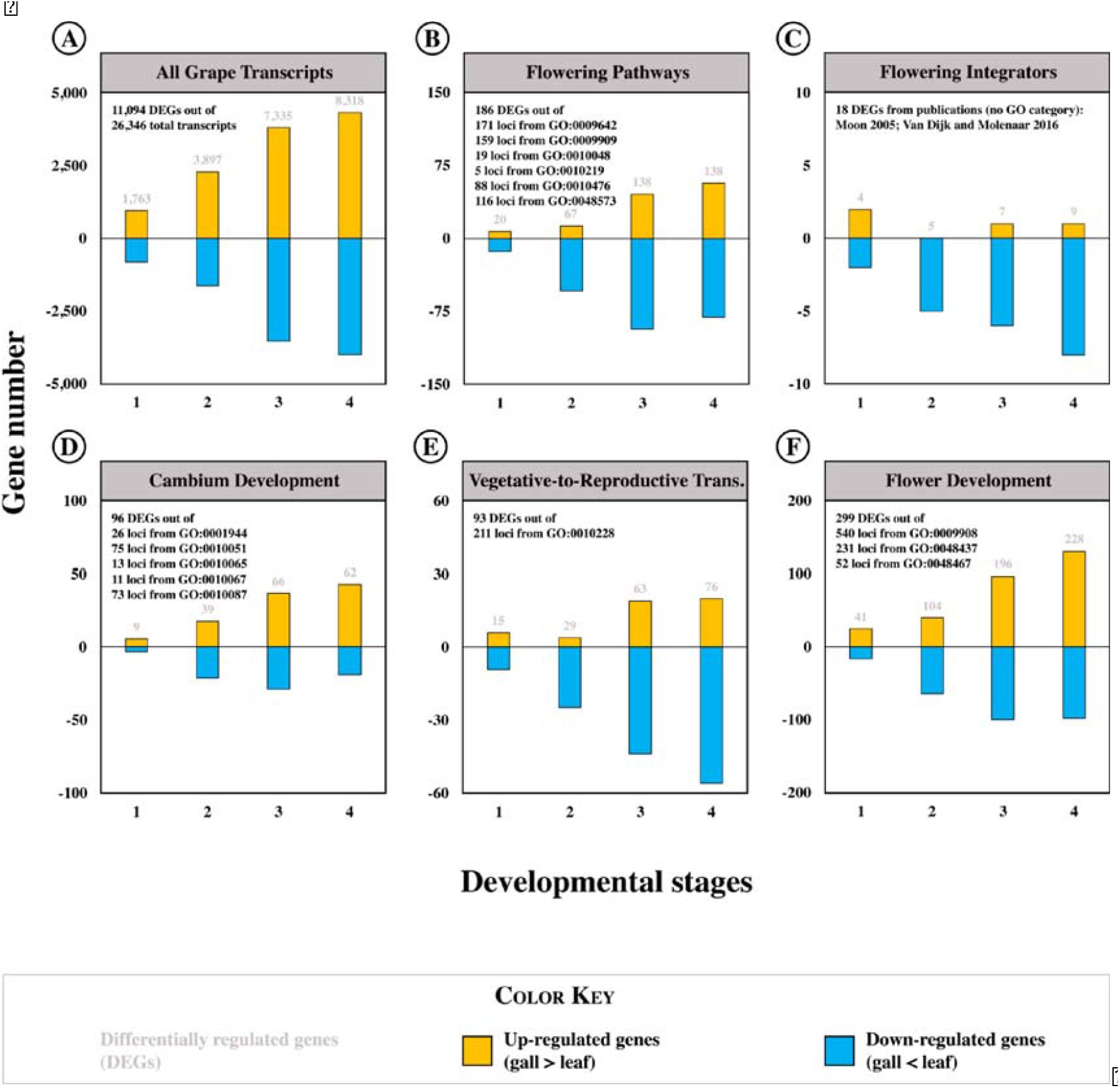
Number of sequences that were significantly differentially expressed in galls compared with leaves, organized by G0 category. (A) The number of all DEGs increases as the galls and leaves develop. (B) The number of DEGs from canonical flowering pathways increase with development, but many are downregulated. (C) Most integrative DEGs are downregulated throughout development. (D) The number of DEGs involved in cambium/meristem development and activation increase with development and are primarily upregulated in galls. (E) The number of DEGs involved in reproductive transition increases with development; many are downregulated. (F) The number of DEGs involved in development of flowers increases with development.

The number of transcripts expressed differentially (DEGs) in galls compared with leaves increased dramatically as the gall developed, from 1,763 in stage 1 to 8,318 in stage 4 (Fig. 2.).

The functional makeup of transcripts in developing galls was distinct from that in age-matched ungalled leaves. We assorted transcripts that were significantly up- or down-regulated in galls compared with leaves into gene ontology (GO) categories using the PANTHER classification system^17^. Significantly enriched GO categories related to reproduction were present throughout gall development but their number increased dramatically in later gall development stages (Table 1). Reproductive GO categories were enriched among both upregulated and downregulated DEGs throughout gall development, although more frequently among upregulated DEGs (Table 1).

**Table 1.** Enrichment of GO categories related to reproduction among DEGs from galls compared with leaves. Values are -fold enrichment.

| GO Term | Search Term | Up-regulated genes |  |  |  | Down-regulated genes |  |  |  |
| --- | --- | --- | --- | --- | --- | --- | --- | --- | --- |
|  |  | Developmental stages |  |  |  | Developmental stages |  |  |  |
|  |  | 1 | 2 | 3 | 4 | 1 | 2 | 3 | 4 |
| GO:0010154 | Fruit development | 2.31 | 0.00 | 1.57 | 1.74 | 0.00 | 0.00 | 1.74 | 0.00 |
| GO:0048316 | Seed development | 2.29 | 1.83 | 1.54 | 1.69 | 0.00 | 0.00 | 1.75 | 1.42 |
| GO:0048608 | Reproductive structure development | 1.86 | 1.46 | 1.45 | 1.56 | 0.00 | 1.71 | 1.71 | 1.48 |
| GO:0003006 | Developmental process involved in reproduction | 1.69 | 1.54 | 1.33 | 1.50 | 0.00 | 0.00 | 1.57 | 1.38 |
| GO:0000003 | Reproduction | 1.51 | 0.00 | 1.28 | 1.50 | 0.00 | 1.47 | 1.49 | 1.30 |
| GO:0048438 | Floral whorl development | 0.00 | 0.00 | 2.21 | 2.15 | 0.00 | 0.00 | 0.00 | 0.00 |
| GO:0048443 | Stamen development | 0.00 | 0.00 | 2.19 | 2.12 | 0.00 | 0.00 | 0.00 | 0.00 |
| GO:0048437 | Floral organ development | 0.00 | 0.00 | 1.84 | 1.89 | 0.00 | 0.00 | 0.00 | 1.89 |
| GO:0009908 | Flower development | 0.00 | 0.00 | 1.55 | 1.64 | 0.00 | 2.31 | 1.69 | 1.51 |
| GO:0090567 | Reproductive shoot system development | 0.00 | 0.00 | 1.53 | 1.63 | 0.00 | 2.36 | 1.69 | 1.51 |
| GO:0048465 | Corolla development | 0.00 | 0.00 | 0.00 | 4.40 | 0.00 | 0.00 | 0.00 | 0.00 |
| GO:0009553 | Embryo sac development | 0.00 | 0.00 | 0.00 | 2.13 | 0.00 | 0.00 | 0.00 | 0.00 |
| GO:0009909 | Regulation of flower development | 0.00 | 0.00 | 0.00 | 0.00 | 0.00 | 2.29 | 0.00 | 0.00 |
| GO:0009911 | Positive regulation of flower development | 0.00 | 0.00 | 0.00 | 0.00 | 0.00 | 4.12 | 0.00 | 0.00 |
| GO:0048573 | Photoperiodism, flowering | 0.00 | 0.00 | 0.00 | 0.00 | 0.00 | 3.76 | 0.00 | 0.00 |
| GO:2000241 | Regulation of reproductive process | 0.00 | 0.00 | 0.00 | 0.00 | 0.00 | 2.02 | 0.00 | 0.00 |

### Flowering pathways

Normal flowering is initiated at the shoot apical meristem (SAM) in response to environmental cues and endogenous signals *via* several major pathways^18,19,20,21^ (Fig. 3.). These may include the photoperiod, light quality/intensity, vernalization, gibberellin, and autonomous pathways. These flowering pathways are largely conserved among herbaceous plant species like *Arabidopsis* but can very somewhat in woody plants^22^. In grapevine, ambient temperature, light intensity, age and gibberellin (GA) are the primary influences on initiating flowering^22^. There is little evidence of photoperiod or vernalization impacts on flowering in grapevine^22^. We identified differentially-expressed genes from these pathways in our dataset by searching gene ontology categories G0:0010476 *gibberellin-mediated signaling pathway*, G0:0009909 *regulation of flower development*, G0:0048573 *photoperiodism, flowering*, G0:0009642 *response to light intensity*, G0:0009909 *regulation of flower development*, G0:0010048 *vernalization response*, G0:0010219 *regulation of vernalization response*, G0:0009909 *regulation of Hower development*.

**FIGURE 3.**
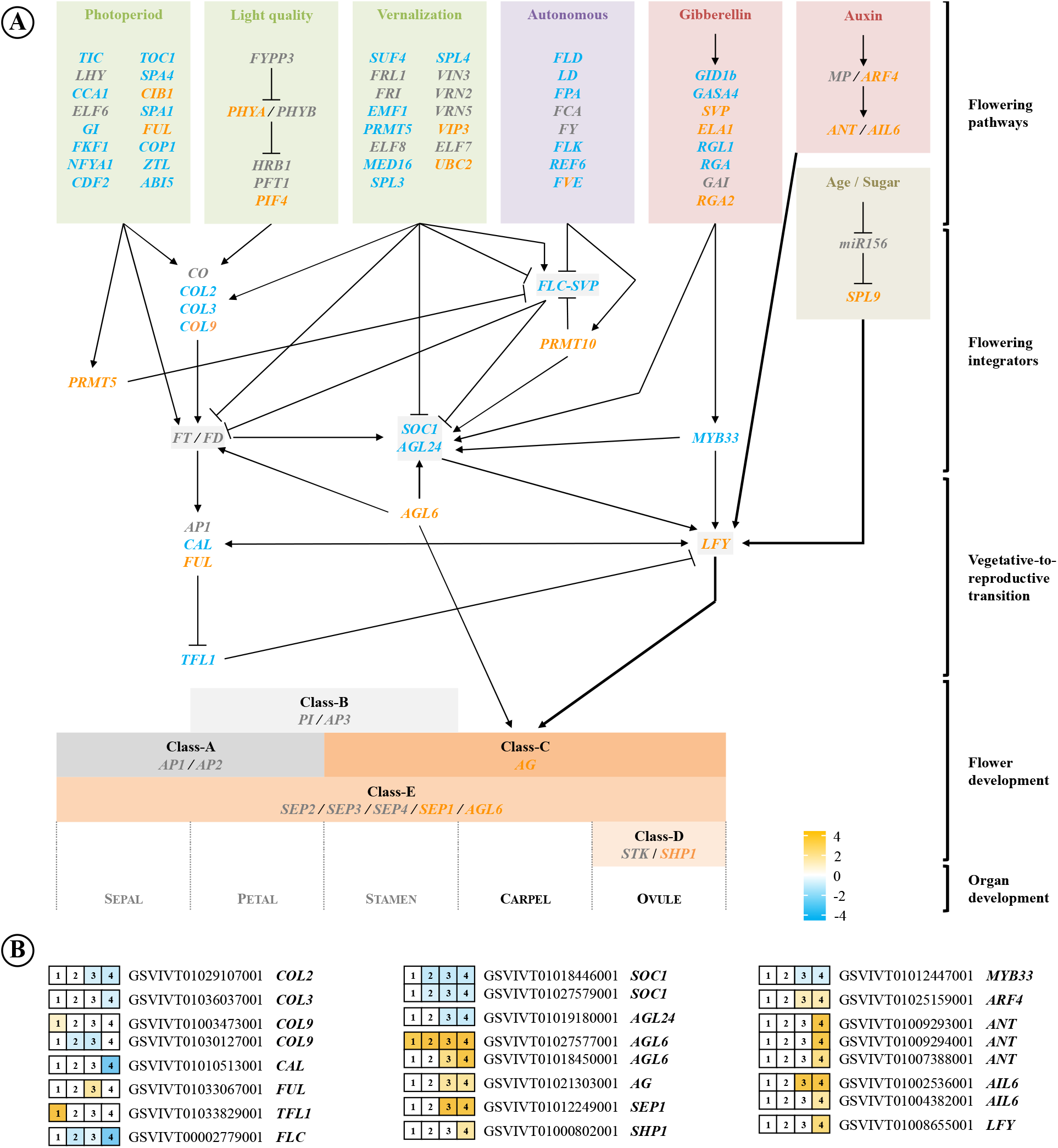
(A) Simplified diagram of key gene pathways regulating flower and fruit development in plants. Activity of canonical flowering pathways is integrated by a few flowering integrators, which regulate the transition from vegetative to reproductive development. Activation of floral meristem identity gene *LFY* promotes flower development via interactions between AG and SEP proteins. The canonical pathways and integrators are blocked in developing galls, while auxin- and age-regulated pathways to *LFY* activation are intact. (B) Expression of selected genes from (A) using RNAseq data. Genes in orange were upregulated in galls, genes in blue were downregulated, and expression of genes in grey (A) or white (B) was unchanged. Dual-color genes had both up- and down-regulated loci.

We identified 162 *Arabidopsis* orthologs of known flowering related genes among 237 *Vitis* loci expressed in phylloxera galls and leaves via RNAseq (Supplementary Figure 3). 0f these, 123 putative genes (184 loci) were differentially expressed in galls. We identified the best-supported function of each DEG ortholog using information curated by TAIR^23^, UNIPROT^24^ and FLOR-ID^25^. We then used this information to infer each DEG’s likely impact on flowering as expressed (up- or down-regulated) in the galls. This examination of the functions of the differentially expressed pathway genes, some of which promote while others delay flowering, revealed that 65 putative genes (91 loci) would promote flowering and 56 DEGs (84 loci) would delay flowering in *Arabidopsis* if they were expressed as they were in galls. About half (80) of flowering pathway genes and 115 loci were differentially expressed only in gall stages 3 and 4 and only 4 genes/loci were differentially expressed exclusively in gall stages 1 and 2 (Supplementary Figure 1).

The particular way in which flowering transition is regulated in grapevine^22^ led us to focus on orthologs related to ambient temperature/light intensity and GA signaling. Two orthologs that may be related to ambient temperature- or light-regulated flowering were differentially-expressed in either the first or second gall stage. Both are normally flowering repressors. One was *ZEITLUPE* (*ZTL*), a flowering suppressor involved in light intensity and photoperiod signaling^26^; it was downregulated. The other was *PHOTOPERIOD-INDEPENDENT EARLY FLOWERING 1 (PIE1)*, which normally suppresses *FLOWERING LOCUS C (FLC)* expression to promote flowering^26^; it too was downregulated. The number of orthologs potentially involved in ambient light or temperature signaling increased through gall development (Supplementary Figure 1). However, each of the DEGs in this category acts by increasing the expression of *FT, SOC1* or G/^6^. The expression of each of these was suppressed or unchanged in galls so that they could not promote flowering there (Fig. 4.).

**FIGURE 4.**
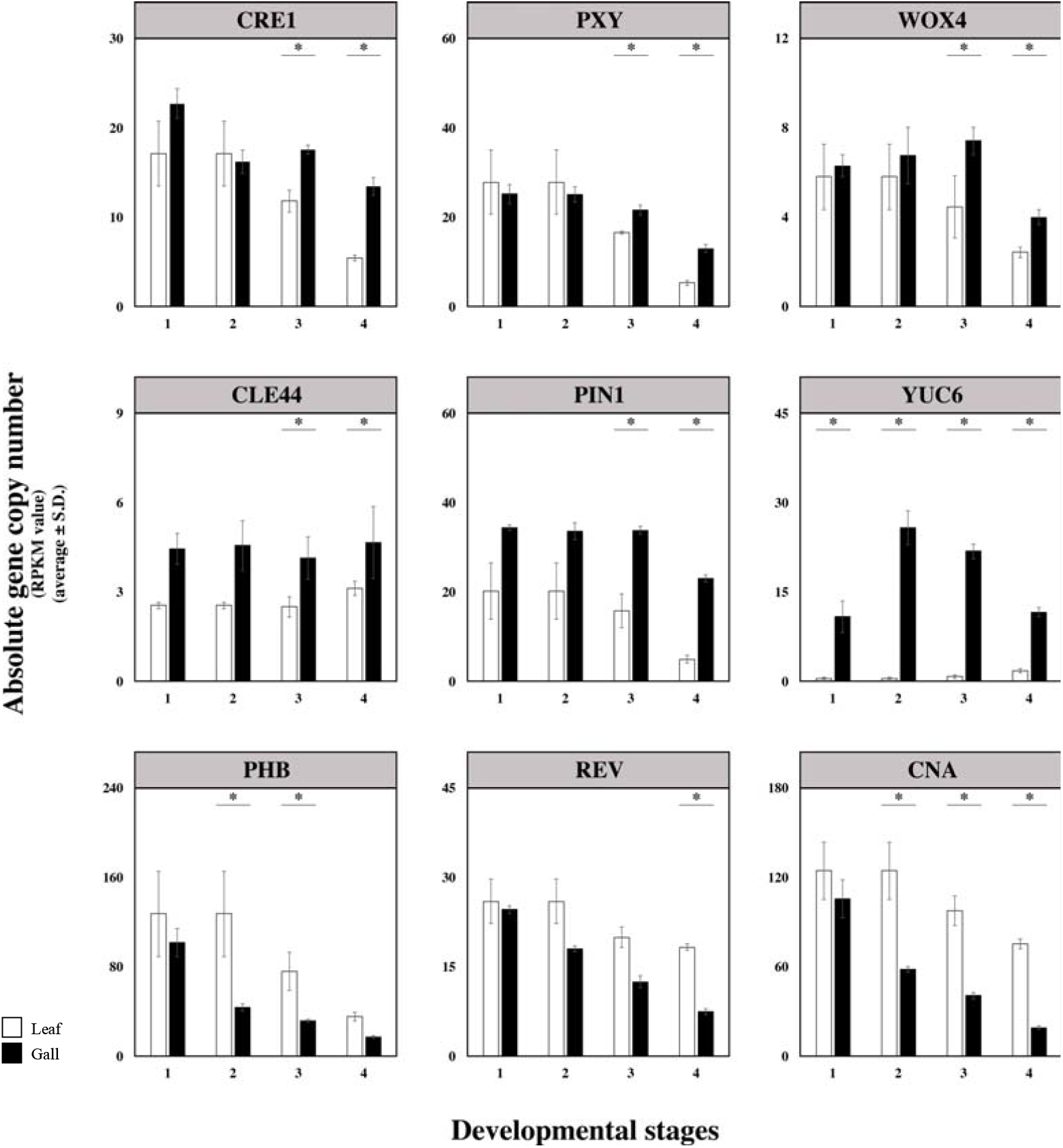
Examples of 3 temporal expression patterns seen among DEGs involved in cambium/meristem activity or development. Top row: DEG expression declines less rapidly than in leaves. Middle row: DEG expression is significantly greater in galls throughout development. Bottom row: DEG expression declines more rapidly in galls than in leaves.

In grapevine, flowering is triggered by an absence or decline in GA signaling^27^ and we observed differences in GA signaling and metabolism as galls developed. Sixteen DEGs (24 loci) from canonical flowering pathways involved in GA biosynthesis or signaling were upregulated in galls, while 26 genes (33 loci) were downregulated in galls (Supplementary Figure 1). Upregulated biosynthesis DEGs included *GIBBERELLIN 20-OXIDASE 1* and *2 (GA20OX1, GA20OX2), GIBBERELLIN 3-OXIDASE 1 (GA3OX1), ENT-KAURENE SYNTHASE (KS)*, and *ENT-COPALYL DIPHOSPHATE SYNTHETASE 1 (CPS1)* (Supplementary Fig 1). Key GA-responsive flowering DEGs included *LFY, AINTEGUMENTA-LIKE 6(AIL6), AGAMOUS-LIKE 6(AGL6), HOMEOBOXGENE 1 (ATH1)*, and *TERMINAL FLOWER 1* (TFL1) (Supplementary Figure 1). Two catabolic *GIBBERELLIN 2-OXIDASE 1* loci *(GA2OX1, GA2OX8)* were downregulated, as was *GIBBERELLIC ACID METHYLTRANSFERASE 2 (GAMT2)*. Key downregulated flowering DEGs included *SOC1, ZTL, FLOWERING LOCUS D (FLD), SHORT VEGETATIVE PHASE (SVP), MYB33, RELATIVE OF EARLY FLOWERING 6 (REF6), FVE, FRIGIDA (FRI)*, and *EARLY FLOWERING 3 (ELF3)* (Fig 3, Supplementary Figure 1).

Since these DEGs may be flowering promoters or repressors, we tallied their likely impact on flowering as expressed. Sixty-two GA-related DEGs would promote flowering in *Arabidopsis*, while 53 DEGs in this list would delay or repress flowering in *Arabidopsis* as expressed in galls. Those numbers reverse if it is indeed true that GA signaling delays or prevents flowering in grapevine^22,27^. Thus, gene expression patterns provide little conclusive evidence about the role of gibberellins in gall development.

### Flowering integrators

The key to initiating flower development is activating the floral meristem initiator *LFY*^28,29^. In normal flowering, these regulatory pathways converge on a small set of floral pathway integrator genes^30,31^. In *Arabidopsis* and other plants, expression of one or more of these integrators must be increased or decreased to allow the floral meristem identity gene *LFY* to initiate flowering^32^. The key integrators are *CONSTANS(CO), SUPPRESSOR OF OVEREXPRESSION OF CO 1 (SOC1), AGAMOUS 24 (AG24), FLOWERING LOCUS T(FT)*, and *FLC. CO* and *FLC* integrate signals from the various pathways and promote or inhibit flowering *via* their impact on expression of *FT* and *SOC1/AGL24*, which in turn activate *LfY*^13,34,35^(Fig. 3.). The influence of the GA pathway is mediated by *SOC1* and the GAMYB transcription factor *MYB3*^6^’^37^.

Four of the flowering integrators we found were upregulated in galls while 11 were downregulated or unchanged in galls (Fig. 2, Supplementary Figure 2). Based on their functions in *Arabidopsis* and grapevine, upregulation of *FRUITFUL (FUL)* and *LFY*, plus downregulation of *MADS AFFECTING FLOWERING 1 (MAF1), CONSTANS-LIKE 3* and *9 (COL3, COL9*, redundant homologs of CO^38^, *CAULIFLORA* (CAL), and *FLC* could all participate in flowering promotion. Downregulation of *SOC1, AGL24*, GAMYB transcription factor *MYB33*, and *COL2* in galls would normally contribute to floral suppression. The expression of *FT* did not differ between galls and leaves and was barely detectable in either; expression of the related *MOTHER OF FT AND TFL (MFT)* was downregulated in galls. Neither expression pattern would support flowering. Downregulation of *FLC* in galls could facilitate flowering, but since its impact on *LFY* and vegetative-reproductive transition depends on the activity of *FT* and *SOC1* - which were unchanged or suppressed - *FLC* is unlikely to permit or promote flowering processes to proceed in galls. *LFY* and *FUL* comprise the only flowering integrators likely to promote flower development in galls.

### Sources of meristem

Flowering is normally initiated at the apical meristem in response to the signaling pathways and integrators described above. Since gall development is a form of *de novo* organogenesis, it presumably requires stem cells as a starting point^39,40^. While some galls appear on apical buds, phylloxera galls and many others form on leaves or stems. In leaves, the only meristem is (pro)cambium from which vascular tissue is derived.

To determine whether cambial meristem might provide a foundation for gall development, we examined the expression of genes in GO categories specific or related to vascular cambium formation and activity: G0:0010067 *procambium histogenesis*, G0:0010065 *primary meristem tissue development*, G0:0010305 *leaf vascular pattern formation*, G0:0001944 *vascular development*, G0:0010087 *phloem orxylem histogenesis*, and G0:001005 *xylem and phloem pattern formation*. We found 96 orthologous loci from these categories differentially expressed in galls; expression of 44 genes (54 loci) was significantly greater in galls than leaves (Fig. 2, Supplementary Figure 3). Most (67 genes, 91 loci) meristem-related activity occurred in stages 3 and 4 on more mature leaves).

Several broad functional groups can be seen in these meristem-related DEGs. Seventeen DEGs (23 loci) are involved in forming, activating, or maintaining vascular cambium Supplementary Figure 3). These include upregulation in galls of the gene encoding a signaling peptide, *CLAVATA3/ESR-RELATED 44 (CLE44)*, its receptor *PHLOEM INTERCALATED WITH XYLEM (PXY)*, PXYtarget *WUSCHEL RELATED HOMEOBOX 4 (WOX4)*, and *ERECTA* (ER), which together form a multifunctional pathway that regulates cambium stem cell pools^41,42^. 0ther DEGs involved in regulating (pro)cambium function including *ETHYLENE RESPONSE FACTOR 104* and *109 (ERF104, ERF109), VEIN PATTERNING (VEP1), CYTOKININ RESPONSE 1 (CRE1), AUXIN RESPONSE FACTORs 3* and *4 (ARF3, ARF4), TARGET OF MONOPTEROS 6 (TMO6), SHRUBBY (SHR), HIGH CAMBIUM ACTIVITY 2 (HCA2), LITTLE ZIPPER 3 (ZPR3)* and *VASCULATURE COMPLEXITY AND CONNECTIVITY* (*VCC)* were upregulated in galls compared with leaves, with the exception of one of the two *ERF2* loci. Ethylene signaling can stimulate cell division in cambium of *Populus*^43^ and *Arabidopsis*^44^.

We identified 22 DEGs (29 loci) involved in more general meristem initiation, maintenance or growth. These included 5 loci of *ALTERED MERISTEM PROGRAM 1* (AMP1), *PENNYWISE (PNY), POUNDFOOLISH(PNF), CLAVATA 1* and 2 *(CLV1, CLV2), CORYNE(CRN), ARGONAUTE10(AGO1Q)* and *REVOLUTA (REV)*. All were upregulated in galls compared with leaves except for *REV* and one locus of *AGO10* (Supplementary Figure 3).

Stem cell state and availability to phylloxera for programming gall development presumably ends when cambium cells differentiate as vascular tissues. We found 9 DEGs (11 loci) involved in vascular differentiation (Supplementary Figure 3). Six of the 11 loci that promote vascular differentiation were downregulated in galls. Examples include *TARGET OFMONOPTEROS 5-LIKE(TMO5-LIKE), DEFECTIVELY ORGANIZED TRIBUTARIES 3*and *4 (DOT3, DOT4), REDUCED WALL ACETYLATION 1 (RWA1), VASCULAR RELATED NAC-DOMAIN PROTEIN 4* (*VND4), CORTICAL MICROTUBULE DISORDERING1 (CORD1), PHABULOSA (PHB)*, and *IRREGULAR XYLEM8 (IRX8)*. The 4 vascular differentiation-related DEGs (5 loci) upregulated in galls negatively regulate vascular differentiation, mainly by extending cambium cell division activity^41^. These include ethylene-response factors *ERF104*and *ERF109, MYB61*, and *ER*.

Twenty DEGs (32 loci) associated with establishing polarity or pattern in vascular development were upregulated in galls. Examples include *KANADI(KAN), TORNADO 1* and *2 (TRN1, TRN2), AMP1, ASYMMETRIC LEAVES 2 (AS2), VEIN PATTERNING 1 (VEP1)*, and *PNY* (Supplementary Figure 3). These genes are involved in specifying the precise location of auxin in developing organs ^45,46,47^

Development, growth and patterning of cambium and the vasculature are regulated by phytohormones. Signaling by or responses to the phytohormone auxin as they relate to cambium activity^45,48^ was indicated by expression of 14 DEGs, including *PIN-FORMED 1 (PIN1), ARF2, 3* and *4, LIKE AUXIN RESISTANT 2 (LAX2), TRN1, VHI-INTERACTING TPR CONTAINING PROTEIN (VIT), AS2, LONESOME HIGHWAY (LHW), DOT3, VASCULAR HIGHWAY 1* (VH1), *REV, PHABULOSA (PHB)*, and *ACAULIS 5(ACL5)*. Also activated in galls were three DEGS involved in auxin synthesis, *TRYPTOPHAN AMINOTRANSFERASE 1 (TAA1)* and *TRYPTOPHAN AMINOTRANSFERASE RELATED 2 (TAR2)*, plus *YUCCA6 (YUC6)*, which controls the formation of vascular tissues as well as floral organs in *Arabidopsis^49^*. Cambium-related cytokinin signaling in galls was suggested by elevated expression of *CYTOKININ RESPONSE 1 (CRE1)*. However, cytokinin activators *LONELY GUY 1* and *3 (LOG1, LOG3)* were downregulated in late stage galls (Supplementary Figure 3).

The divergence in expression of cambium-related genes in gall and leaf as they developed exhibited several different temporal patterns (Fig. 4.). Expression of many genes declined in both leaves and galls as they aged, but less rapidly in galls, producing statistically significant differences by gall stage 4 (Fig. 4.). In a second pattern, gall values showed little or no decline with development and more or less exceeded leaf values over the entire course of development (Fig 4). A third pattern involved gall values that declined more precipitously than values in leaves (Fig. 4.).

### Vegetative-to-reproductive transition

*LFY* is the master initiator of floral meristem development and indicator of the vegetative-reproductive meristem transition^29,49^. *VFL*, the grape homolog of *AtLFY*, functions similarly in the grapevine flowering transition^50^. Having established that the canonical flowering pathways appear unlikely to trigger elements of flower development in galls, we examined expression of flowering triggers by identifying DEGs in our gall data set found in the G0 category *vegetative to reproductive phase transition of meristem* (G0:0010228). We found altered expression of 76 genes (93 loci) from that G0 category in developing galls (Fig. 2; Supplementary Figure 4). Thirty-nine (43 loci) would promote the transition to flowering in *Arabidopsis* if they were expressed as they were in galls, while 30 DEGs (42 loci) would repress it.

While all 76 genes were differentially expressed in the 3^rd^ and 4^th^ gall stages, several genes involved in the vegetative-to-reproductive transition were also expressed in the earliest stages (Supplementary Figure 4). These included *AGL6, PROTEIN ARGININE METHYLTRANSFERASE10 (PRMT10)*, one locus of *COL9*, and *GA20OX1*, all of which were upregulated in the youngest galls. Genes that were downregulated early include *SUPPRESSOR OFPHYA-1051 (SPA1)* and *DNAJHOMOLOGUE 3(J3)* (Supplementary Figure 4). However, these and many other DEGs in this category act as part of, or together with, one or more canonical flowering pathways and depend for their influence on the flowering integrators we found inactive or downregulated (Fig. 3.).

The same vegetative-reproductive transition DEG set included meristem transition triggers not affiliated with the canonical pathways and their integrators. Krizek^51^ and Yamaguchi et al.^29^ have described an auxin-responsive pathway in *Arabidopsis* leading to flowering, dependent on *AINTEGUMENTA (ANT), AIL6* and *LFY* (Fig. 3.). They showed that *ANT* and *AIL6* expression is elevated in response to auxin, and that they in turn activate *LFY* to initiate flowering. Auxin sources include polar transport involving *PIN1* as well as synthesis by members of the *YUCCA (YUC)* family^51^; expression of both was elevated in developing galls (Fig. 3, Supplementary Figure 4). Krizek^51^ implicated auxin response factors ARF3 and ARF 4 in this signaling network. We found elevated expression of *ARF2, ARF3, ARF4* and *ARF6* orthologs in developing galls (Fig. 3, Supplementary Figure 4). In *Arabidopsis ARF4* is a target of *LFY^52^* and regulates polarity^53^, *ARF6* regulates gynoecium maturation and *ARF2* and *ARF3* are involved in carpel and ovule development^54,55^. All of the elements of auxin-triggered transition to flowering were activated in developing galls (Fig. 3.).

An age-based pathway to flowering transition was also active in developing galls (Fig. 3.). Plants must mature over some period of time before they become competent to flower^56^. Grapevine generally requires 3-6 years before it can reproduce^57^. As plants age, the expression of micro RNA miRNA156 decreases. miRNA156 suppresses expression of the transcription factor *SQUAMOSA PROMOTER BINDING-LIKE 9 (SPL9)*, which is a promoter of *LFY* expression. As miRNA156 activity decreases, *SPL9* expression increases and eventually increased *LFY* expression triggers flowering, independent of the canonical flowering pathways. While we could not assess miRNA abundance or activity using our methods, the expression of *SPL9* increased significantly in galls as they aged; this increase could promote the flowering process in galls.

Some gall DEGs found in G0:0010228 influence the flowering transition *via* pathways or genes that were not found to be activated in galls. For example, *AGL6, PRMT10, PRMT5, J3, SQUAMOSA PROMOTER BINDING PROTEIN-LIKE 3 (SPL3)*, and *REF6* all influence the transition to flowering by elevating expression of *FT* or SOC1(58), neither of which was activated in galls (Fig. 3, Supplementary Figure 3).

### Flower development

To determine the degree to which genes that direct actual floral organ development might be involved in gall development, we examined the expression of DEGs in the gene ontology categories *floral organ development* (G0:0048437) and *flower development* (G0:0009908) (Supplementary Figure 5). We identified 227 putative ortholog genes (296 loci) from those two categories differentially expressed in developing galls (Supplementary Figure 5). 0f these, 118 DEGs (154 loci) were upregulated in galls compared with leaves and 121 (142 loci) were downregulated in galls. After identifying roles in flower development, we found that 142 DEGs (181 loci) would promote development of floral organs in *Arabidopsis* as expressed in galls while 87 DEGs (105 loci) would repress or not affect flower development (Supplementary Figure 5).

While *LFY* is the master regulator and indicator of floral meristem development, it also triggers the transcription of key components of flower organ determination through its interaction with *AGAMOUS* (AG)^59^. We found that *LFY* expression was significantly elevated in gall stage 4 (Table 1, Fig. 3, Supplementary Figure 5), whereas its target AG, which terminates meristem activity so that floral organogenesis can proceed^59^, was significantly upregulated in gall stages 3 and 4 (Supplementary Figure 5). This chain of events is normally repressed by *TERMINAL FLOWER (TFL)* in both *Arabidopsis* and grapevine^60^. Expression of the ortholog of the *Arabidopsis TFL* was upregulated during gall stage 1, but subsequently declined to leaf levels as galls developed (Supplementary Figure 5). *MOTHER OF FT AND TFL (MFT)*, which functions similarly in grapevine^60^, was downregulated in gall stages (Supplementary Figure 5). Altogether, we found 22 DEGs (27 loci) involved in the decision to maintain floral meristems or allow differentiation to proceed (Supplementary Figure 5). The majority, 18 DEGs (23 loci), would lead to floral differentiation in *Arabidopsis* if expressed as in phylloxera galls. 0f the 4 DEGs that do not directly promote floral meristem activity, one (*STM*) requires the combined activities of *FT* and *SOC1*, which were not differentially expressed (Fig 3). Another, *LATE MERISTEM IDENTITY2 (LMI2)*, was downregulated in galls. It interacts with *LFY* but is not necessary for flower formation^61^. Upregulated *REBELOTE* (RBL) contributes to floral meristem termination so as to prevent the formation of supernumerary flowers or floral organs^62^. The activation of *LFY* and *AG* in developing galls should set the stage for flower organ development.

To determine how carpel development and related genes might be involved in gall formation, we examined the expression of all unique genes from ontology category *gynoecium development* (G0:0048467) augmented with a list developed by Reyes-0lalde^63^ (Supplementary Figure 5). We found expression of 39 orthologs (39 loci) to be elevated in galls compared with age-matched leaves. These include^63^ *NO TRANSMITTING TRACT* (NTT), *SEPALLATA 1 (SEP1), ASYMMETRIC LEAVES 2 (AS2), ASYMMETRIC LEAVES 2-LIKE 1 (ASL1), JAGGED (JAG), PERIANTHIA (PAN), PHABULOSA (PHB), YABBY 1 (YAB1), NGATHA1 (NGA1), SHORT VALVE1 (STV1), SHATTERPROOF2(SHP2), AGAMOUS(AG), FRUITFULL (FUL), ULTRAPETALA1 (ULT1), AINTEGUMENTA (ANT), AIL6, WUSCHEL RELATED HOMEOBOX 13(WOX13), SPATULA (SPT)*, and *HECATE 1 (HEC1)*, among others (Supplementary Figure 5). All of these genes would participate in carpel/gynoecium development in *Arabidopsis* if expressed as they were in galls. At the same time, carpel development repressors *SHORT VEGETATIVE PHASE (SVP), LEUNIG (LEU), EARLY FLOWERING IN SHORT DAYS (EFS)* were downregulated (Supplementary Figure 5). *AGAMOUS* repressors *SEUSS, PAN, FLC*, and *BELL-LIKE 1* (BEL1)^64,65,66^ were also downregulated in galls(Supplementary Figure 5).

Carpel/gynoecium development is regulated by phytohormones, and G0:0048467 includes phytohormone-related genes. Phytohormone activity in stage 4 galls was indicated by upregulation of gynoecium development genes *CYTOKININ OXIDASE 3* and *5 (CKX3, CKX5), TAA1, TAR2, ARF2, ARF3, ARF6, PINOID (PID), PIN1, BRASSINAZOLE-RESISTANT 1* (BZR1), BRASSINAZOLE-INSENSITIVE1 (BIN1) and *BRASSINOSTEROID-6-OXIDASE 2 (BR6OX2)* (Supplementary Figure 5).

Once the vegetative-to-reproductive transition has been achieved, *AG* interacts with floral homeotic genes to regulate floral organ development in *Arabidopsis* and other species^67^ (Fig 3). Proteins encoded by a small number of homeotic genes interact in a combinatorial way to determine each of the major floral organs: sepals, petals, stamens, and carpel^67^. The homeotic genes required to produce these structures have been classified A, B, C, D, or E^67^. We found no differential expression of orthologous homeotic genes from class-A or −B (Fig. 3, Supplementary Figure 5). However, orthologs of the class-C carpel identity genes *AG^6^* and *SHATTERPROOF 1* (SHP1)^69,70^ were strongly upregulated in gall stages 3 and 4 compared with leaves (Supplementary Figure 5). Class-C proteins interact with class-E proteins to direct development of the floral organs^71^ (Fig. 3.). In *Arabidopsis* the major class-E genes comprise the *SEPALLATA* family^72^. The combination of AG and SEPx is required to produce a carpel^68^. *SEPALLATA 1 (SEP1)* was strongly upregulated in galls (Supplementary Figure 5). The protein encoded by *AGL6*, which was strongly upregulated throughout gall development, also fulfills SEPx functions in some plant species^73^. All the elements necessary for flower development, from activated *LFY* through *AG* expression to elevated transcripts for *SEP1* and *AGL6* are present in phylloxera galls.

## Discussion

We found that gall and leaf transcriptomes differ at the earliest point in gall development, and diverge increasingly as galls and leaves develop. The transcription of many grape genes orthologous or homologous to genes responsible for triggering flowering and regulating flower development in *Arabidopsis* is altered in phylloxera leaf galls. The general pattern is that expression of these genes, many of which have little or no role in the development of the leaf on which the gall grows, is up- or down-regulated in ways that could lead to flowering and eventual fruiting. Expression of many floral repressors were found to be downregulated, while promoters were upregulated. The frequency of differentially-expressed flowering genes increased dramatically as the gall developed and the leaf matured.

Flowering requires a transition from a vegetative state to the reproductive meristematic state. This transition is elicited by the influence of environmental or hormonal signals on a few key floral integrator genes, which in turn increase the expression of the master regulator *LFY* to establish a floral meristem and promote flower development^32^. Indeed, ectopic *LFY* expression is sufficient to produce flowers in the absence of repressors^74^. 0verexpressing these genes as well as other flowering genes in *Arabidopsis* can produce ectopic flower development, particularly carpels, or gall-like morphological changes ^49,71,75^*LFY* expression was significantly elevated in late-stage galls. We therefore consider the expression of *LFY* and its targets a key step if gall development involves aspects of flower development and so investigated all of the ways in which *LFY* expression could be elicited.

### Flowering pathways

We first asked whether phylloxera could be exploiting the canonical pathways that culminate in activating *LFY* to trigger flowering (Fig. 3.). We found that the differences between galls and leaves in the expression of the many grape orthologs of *Arabidopsis* genes in those pathways were mixed. Some pathway orthologs were expressed in ways that would prevent their impact on *LFY* while expression of others could promote *LFY* expression.

Expression of genes in the gibberellin pathway was consistent with GA’s role in normal flower promotion in many plant species. For example, orthologs of many GA biosynthesis and response genes were upregulated in late gall stages while catabolic genes were downregulated (Fig. 3, Supplementary Figure 1). However, GA signaling suppresses flowering in grapevine^27^ so positive GA signaling could prevent flower development as part of gall development. The impact of GA signaling on flower development may depend on signaling by the GAMYB transcription factor MYB33 ^36,37,76^ *MYB33*was downregulated in late-stage galls, in principal blocking GA signaling. Since gibberellin’s influence on flowering switches from negative to positive during flower development^77^, a more detailed study of the timing of GA signaling will be needed to determine its role in gall development.

Overall, we did not find convincing evidence that gall elicitation or development depends on the canonical flowering pathways as they normally function in flowering.

### Flowering integrators

Signaling by all the canonical flowering elicitation pathways converges on a few integrating genes^18,58^. These integrators in turn elevate *LFY* expression to bring about the meristem transition to flowering and flower development^49^. The only integrator gene expressed in galls in a way that would influence *LFY* expression was the floral repressor *FLC*. However, *FLC* s impact on flowering comes about when it is downregulated and its repression of *FT* and *SOC1/AGL24* is stopped. Expression of *FT* and *SOC1/AGL24* was unchanged or decreased in galls as compared with leaves. This fact alone would appear to rule out most or all canonical environmental signaling pathways as gall elicitors, (Fig. 3.).

### Vegetative-to-reproductive transition

Indicators of a meristematic transition from vegetative to reproductive state were conspicuous in galls. Genes involved in floral meristem identity and/or maintenance were upregulated in gall stages 3 and/or 4, including *LFY*, *AG*, *FUL*, *CAL*, and *UNUSUAL FLORAL ORGANS (UFO)*. 0ne exception was *AP1*, which was unchanged. *AP1* expression is not associated with flowering in grapevine^78^. *AG* has a dual role in floral meristem identity early and meristem termination plus organ differentiation later^79^. *TFL* and relatives, which repress floral meristem formation, were unchanged or downregulated in late gall developmental stages.

We found evidence suggesting that localized auxin signaling could play a role in the vegetative-reproductive transition and gall development. Auxin signaling mediated by auxin response factors (ARFs) and acting *via* expression of *ANT* and *AIL6* can elevate *LFY* expression and lead to flowering transition and flower development in *Arabidopsi*s^29^. All of the orthologs in this short pathway were significantly elevated in late gall stages, as were other auxin-responsive signaling and biosynthetic genes (Fig. 3.). Phylloxera could initiate flowering processes *via* local elevation of auxin concentrations or signaling.

An age-related flowering pathway could also be involved in gall development. Like most woody plants, juvenile grapevines require a maturation period of several years before becoming reproductively competent. During this time the expression of microRNA miR159 declines, and its suppression of *SPL9* decreases. Increasing expression of *SPL9* then provokes *LFY* expression to trigger flowering^56^. *SPL9* expression was significantly elevated about 2-fold in late stage galls, but our methods provided no evidence concerning miR159 expression. Medina *et al*.^80^ found that miR159 played a role development of galls elicited by the root-knot nematode *Meloidogyne incognita*. The potential role of microRNAs in insect gall development warrants further attention.

### Sources of meristem

As in normal flowering and organogenesis, gall development requires undifferentiated stem cells. Normal flowering is initiated at the SAM in response to hormonal and/or environmental cues. There is no SAM in plant leaves, but we found evidence that vascular cambial meristem remains active in galls long after it declines in the leaf and so is a possible source of stem cells for exploitation by the insect. Phylloxera galls (and many others) are always associated with leaf veins and may obtain undifferentiated cells there from which to develop a novel organ. Expression of the key cambial activation genes, *WOX4, CLE44*, and *PXY*^41,42^ was significantly elevated in galls compared with leaves as leaves and galls aged. We found elevated expression of genes associated with hormonal signaling normally involved in cambium activation and reduced expression of genes that terminate cambium activity and promote vascular differentiation well after leaves and their vasculature were mature. While activated cambium could reflect increased vascular development, phylloxera galls do not exhibit increased vascularization^81^. The gall transcriptome is consistent with phylloxera manipulating vascular cambium to provide stems cells for organ development.

Expression of the *WOX, CLE*, and *PXY* cambial activation pathway is also key to development of the gall-like structures elicited by root-knot and cyst nematodes and nodulation by *Rhizobium* in legumes^82^, suggesting that phylloxera and other parasites have converged on altering developmental regulation of vascular stem cells during gall elicitation. We are aware of no studies of CLE peptide production in phylloxera or other insects, as has been shown to be important for root-galling nematodes^82^.

### Flower development

We found transcriptional indications of flower development, including organ determination, in the phylloxera galls. Many orthologs of genes that positively regulate flower development were upregulated. Differential expression of grape orthologs of canonical *Arabidopsis* “ABCE” model homeotic genes that determine flora organ identities was significant for class-C genes. Class-C AG protein normally associates with class-E SEP proteins to determine carpel identity^83^, and expression of *Vitis AG, SHP*and *SEP1* orthologs was strongly upregulated in galls. We also found enhanced expression of the grape ortholog of *AGL6*, a close relative that plays a *SEP* role in several other species^73,84^ and has an ancestral role in carpel identity^79^. Expression of *HUA1*, a regulator of stamen and carpel identities in *Arabidopsis*^85^ as well as other carpel/gynoecium identity genes, was also elevated in galls. The phylloxera gall resembles a carpel more than any other floral organ transcriptionally, anatomically and functionally.

The view that galls are convergent on carpels or fruit is supported by diverse observations from other studies. At least one gall’s nutritive layer includes proteins normally found only in the seed^86^. Gall development and growth of the nutritive layer depend on chemical cues from the insect^87^, much as embryos direct development of surrounding tissues hormonally^13^. At least one galling insect invades fruits and takes over control of normal endosperm development in the absence of the plant embryo^88^. Defensive chemistry is sequestered away from the insect in the gall as it is from embryos in fruits^2,12^. Rolled leaf edges are considered ancestral elements in the evolution of some gall lineages and may also be a key innovation in the origin of the carpel^3,89^.

The absence of evidence for signaling from the canonical flowering pathways led us to examine other means by which flowering can be elicited. There are many paths to flowering, some of which are independent of environmentally-cued pathways. All of the known pathways generally culminate in hormone-regulated gene expression. Most of the major plant hormones have been found to play some role in flowering, and their interaction during flowering and flower development is complex. 0ur results and these observations suggest that direct provision or manipulation of phytohormones is the most plausible means of gall elicitation, although we cannot rule out the injection of CLE peptides or small RNAs, which has not been described in insects.

The idea that galling insects somehow manipulate plant hormones to accomplish their ends is very old, and accumulation of various hormones in galls has been reported frequently^90^. *LFY* responds to both GA and auxin^91,92^. Manipulating signaling by one or more of these hormones would seem a likely way for galling insects to trigger flowering programs in producing a gall. 0ur results, which found elevated expression of auxin-responsive genes and auxin transport genes in the galls, suggest an important role for auxin in phylloxera gall formation. The requirement for local auxin accumulation to prompt organ development, including flowers, is well established^51,91^. 0n the other hand, our results suggest that gibberellin signaling may be suppressed in developing galls, which could stimulate reproductive development at gall sites in grapevine^27^. Definitive resolution of hormone signaling in gall development will require an integration of detailed chemical and transcriptional analyses.

### Limitations to this study

Our conclusions are based on the assumption that similarity between computed *Arabidopsis* and *Vitis* protein sequences suitably indicates similarity in function for a given gene in both *Vitis* and *Arabidopsis*. While we are confident in the assignment of orthologs between the two species, this assumption about functional similarity is no doubt more valid for some genes than for others due to expansion of some gene families in *Vitis* and sub- or neo-functionalization. Network level rewiring may have altered activator and repressor roles of transcriptional regulators in *Vitis* compared to *Arabidopsi^(93)^*. Thus, even when gene families are of similar size, there is no guarantee of a one-to-one function concordance. For example, the key floral meristem gene *LFY* is expressed in a wider range of situations in grapevine than is the *Arabidopsis* LFY^50^. While floral meristem indicator *AP1* is key to the development of flowering competence in *Arabidopsis*, that is not true in *Vitis*, where its impact is restricted to tendrils^94,78^. 0n the other hand, the functions and expression of many of the reproductive genes we identified in galls, such as *AG*, *SHP*, the hormone signaling elements, the pathway integrators, and others are highly conserved among plant species and exhibit the same or similar expression patterns in grapevine_18_

We also did not identify putative orthologs for all genes using the current methods. While we might find more matches through broader searches, we are missing very few important flowering genes and none that would significantly change our conclusions. 0ur conclusions are based entirely on transcriptional data, and ignore post-transcriptional and other regulatory mechanisms. In particular our methods did not allow an assessment of the impact of small RNAs, which are important regulators of many reproductive genes including those we studied^31^.

Our conclusions are also based on statistically significant differences in the numbers of RNA transcripts between gall and leaf tissue. Very few genes were present in one tissue and not the other, despite the fact that many are involved in flower development. It is important to remember that flowers are modified leaves, evolutionarily^95^. Phylloxera galls are not flowers or fruits, but their transcriptomes show greater commitment to flowering than do ungalled leaf tissues; they are neither flowers nor leaves, but are unique organs incorporating traits of both.

In summary, we have shown that phenotypic similarities between galls and fruits extend to their transcriptomes. The likely reason for this is that the plant embryo and galling insect have similar requirements for success and manipulate plant development similarly to achieve similar goals. Both need the conditions provided by an expanded carpel. The patterns we obtained support the hypothesis that the phylloxera leaf gall - and probably other similar galls - is developmentally and transcriptionally convergent on floral organs, particularly the carpel.

## Methods

### Study system

Grape phylloxera *(Daktulosphaira vitifolia (Fitch 1855)* is an aphid relative, native to North America, that feeds on leaves and roots of certain *Vitis* species. It elicits complex galls on abaxial leaf surfaces, and causes swelling on roots when feeding there. Its life history and gall development have been described by Sterling^96^ (Fig. 1.). Females emerge from eggs in the spring and feed from the upper surfaces of the youngest leaves, sucking contents from parenchyma cells beneath them. Within 24-48 hours a disk-shaped depression forms under the feeding insect. Cell division and expansion are altered at the disc margins and soon a circular ridge or wall surrounds the feeding insect. Within 48-72 hours the abaxial depression containing the insect deepens due to differential cell division and expansion and the adaxial wall closes over her, leaving a narrow opening protected by dense trichomes. Two tissue layers several cells thick underlie the depression, an inner layer that is densely cytoplasmic and an outer layer that contains larger vacuoles, enlarged nuclei and nucleoli, and cytoplasmic globules. These ‘secretory’ characteristics spread to other cell layers, becoming a thick ‘nutritive zone’. Development of a complete gall takes 4-5 days, at which point the insect has matured and begins producing eggs. “Crawlers” hatch from eggs in the gall, exit through the abaxial opening, and proceed to feed and form galls on younger leaves. Gall development stops if the insect is removed before this final stage.

### Tissue sampling

Galled and ungalled leaves were collected between 0900 and 1000 from April to August 2014 and 2015 from wild *Vitis riparia* Michx. vines near Rocheport, Missouri, USA (38° 58’ 16.424” N, 92° 32’ 54.118” W). Galls from 3 different vines were separated by size into 4 developmental categories^96^ (Fig. 1.) and dissected on ice; midribs were removed from ungalled control leaves. Because the two earliest gall stages developed on the same leaves, there were only 3 control leaf size classes matched to the 4 gall stages. To obtain enough RNA, samples were pooled from 3 individual vines, producing 3 biological replicates for all analyses. All tissues were immediately frozen in liquid nitrogen and stored at −80 °C.

### RNA Extraction

RNA was extracted and DNase1-treated, on column, using the Spectrum Plant Total RNA Kit (Sigma #STRN50-1KT; protocol A and Appendix). The resulting RNA was further purified and concentrated with the RNeasy MinElute Cleanup Kit (Qiagen #74204) and eluted with water. The quality of the resulting RNA was assessed using the Agilent 2100 BioAnalyzer (Agilent, Santa Clara, CA, USA), and all RNA integrity number values were found to be above 8.

### Illumina Library and Construction

The Illumina libraries were constructed using the RNA TruSeq Kit (Illumina, Inc., San Diego, CA, USA), barcoded (TACT ungalled; GTAT galled), and sequenced single-end with 100bp reads on the Illumina HiSeq-2000 platform at the University of Missouri DNA core (HYPERLINK “http://dnacore.missouri.edu”; University of Missouri, Columbia, MO, USA).

### Illumina read processing and expression quantification

A custom Perl script was used to parse the libraries and remove barcode sequences resulting in approximately 40.9 million reads for the ungalled library and 40.3 million reads for the galled library. NextGENe V2.3.3.1 (SoftGenetics, LLC., State College, PA, USA) was used to quality filter the fastq data, remove reads with a median quality score of less than 22, trim reads at positions that had 3 consecutive bases with a quality score of less than 20, and remove any trimmed reads with a total length less than 40bp. Gene expression was quantified using TopHat/Cufflinks software^97^.

Differential expression between galled and ungalled leaf tissue was analyzed for each mapping, using two discrete probability distribution based methods, DESeq and edgeR (HYPERLINK "https://bioconductor.org") and the annotated *Vitis vinifera* V2 genome (DOE-JGI; ftp://ftp.jgi-psf.org/pub/compgen/phytozome/v9.0/Vvinifera/). Read counts and RPKM values (reads per kilobase per million) were calculated for each library. An RPKM cutoff of 0.1 per gene model was applied for comparing expression values. Functional analyses were limited to genes with a differential expression significance <0.05 and >1.5-fold difference.

Genome-wide syntenic analyses were performed to identify *Arabidopsis thaliana* - *Vitis vinifera* orthologs using CoGe HYPERLINK “http://genomevolution.org/CoGe/". In addition, *Arabidopsis - Vitis* orthologs were identified using reciprocal BLASTp analyses (protein databases) with a 0.00001 p-value cutoff resulting in the annotation of ~86.7% of all coding sequences in the *Vitis vinifera* V2 genome.

Gene 0ntology (G0) enrichment analyses were performed for each of the gall and leaf gene expression sets using the PANTHER classification system^17^. Statistical significance for enrichment scores was set at <0.005.

### Validation of RNAseq results with droplet digitalPCR

Purified RNA was converted to cDNA (RT-PCR) with SuperScript III First-Strand Synthesis SuperMix (Invitrogen #11752-050; Invitrogen, Carlsbad, CA, USA). Primers were designed with PrimerSelect (DNAStar, v.13.0.0; DNAStar, Madison, WI, USA) using published *V vinifera* sequences and our own *V. riparia* RNAseq data (Supplementary Table 1). PCR reaction parameters were optimized with qPCR using a MJ Research 0pticon2 PCR thermal cycler (Bio-Rad, Hercules, CA, USA), with iQSYBR Green Supermix (Bio-Rad #170-8882). Droplet digital PCR (ddPCR) reactions were performed on the Bio-Rad QX100 ddPCR System using using QX200TM ddPCR™ EvaGreen Supermix (Bio-Rad #1864034; Bio-Rad, Hercules, CA, USA). Primer sequences, cDNA dilution and volume, and annealing temperature for each gene tested by ddPCR are presented in Supplementary Table 1. Six biological replicates per each of eleven tissue types were used for ddPCR analysis.

We used two abundantly expressed neutral genes whose expression was uniform across all gall and leaf samples, orthologs of *AtDEC* and *AtDNAJ*, as internal controls to normalize the amount of starting RNA used for RT-PCR for all samples (N = 6 biological replicates per developmental stage for both galls and ungalled control leaves). Normalized gene copies for each gene were calculated by dividing their absolute gene copies by the average gene copies of the two neutral genes. Fold-change between galls and their respective ungalled control leaves was calculated for each gene by subtracting the base-2 logarithm of the RPKM value of galls to the base-2 logarithm of the RPKM value of ungalled control leaves followed by one-way ANOVA. ddPCR results for these genes were consistent with results obtained *via* RNAseq (Supplementary Figure 6).

## Data availability

RNAseq data that were generated for this study are available at NCBI Gene Expression Omnibus (HYPERLINK “https://www.ncbi.nlm.nih.gov/geo/”) under study accession XX. The authors declare that all other data supporting the findings of this study are available within the article and its Supplementary Information files, or are available from the authors upon request.

## Acknowledgements

This project was supported by NSF IOS grant #1757358 (JCS HMA). We thank Wade Dismukes and the University of Missouri Informatics Research Core Facility for help analyzing and organizing data; Dean E. Bergstrom for fieldwork, RNA extraction, RTPCR, and digital droplet PCR; J. Chris Pires and Graham N. Stone for helpful discussions; and Karla A. Carter for assistance preparing the manuscript. Special thanks to Bill Spollen for above-and-beyond informatics assistance.

## References

1. Meyer, J. Plant galls and gall inducers (Gebruder Borntraeger, Berlin, 1987).

2. Nyman, T. & Julkunen-Tiitto, R. Manipulation of the phenolic chemistry of willows by gall-inducing sawflies. PNAS 97, 13184–13187 (2000).

3. Rohfritsch, O. Patterns in gall development in Biology of insect-induced galls (eds. Shorthouse, J. D. & Rohfritsch, O.) 60–86 (Oxford University Press, Oxford, England, 1992).

4. Stone, G. G & Cook, J. M. The structure of cynipid oak galls: Patterns in the evolution of an extended phenotype. Proc. R. Soc. Lond. B Biol. Sci. 265, 979–988 (1998).

5. Harper, L. J., Schonrogge, K., Lim, K. Y., Francis, P. & Lichtenstein, C. P. Cynipid galls: Insect induced modifications of plant development create novel plant organs. Plant Cell Environ. 27, 327–335 (2004).

6. Hori, K. Insect secretions and their effect on plant growth, with special reference to hemipterans in Biology of insect-induced galls (eds. Shorthouse J. D. & Rohfritsch, O.) 157–170 (Oxford University Press, New York, 1992).

7. Leitch, I. J. Induction and development of the bean gall caused by Pontania proxima. In Plant galls: Organisms, interactions, populations (ed. Williams, M. A. J.) 283–312 (Clarendon Press, Oxford, 1994).

8. Cook, L. G. & Gullan, P. J. The gall-inducing habit has evolved multiple times among the eriococcid scale insects(Sternorrhyncha: Coccoidea: Eriococcidae). Biol. J. Linnean Soc. 83, 441–452 (2004).

9. Darwin, C. The variation of animals and plants under domestication (ed. Judd, O.) Volume 2 (1868).

10. Larson, K. C. & Whitham, T. G. Manipulation of food resources by a gall-forming aphid: The physiology of sink-source interactions. Oecol. 88, 15–21 (1991).

11. DRehill, B. J. & Schultz, J. C. Enhanced invertase activities in the galls of *Hormaphis hamamelidis*. J. Chem. Ecol. 29, 2703–2720 (2003).

12. Allison, S. D. & Schultz, J. C. Biochemical responses of chestnut oak to a galling cynipid. J. Chem. Ecol. 31, 151–166 (2005).

13. Gillaspy, G., Ben-David, H. & Gruissem, W. Fruits: A developmental perspective. Plant Cell 5, 1439–1451 (1993).

14. Stern, D. L. Phylogenetic evidence that aphids, rather than plants, determine gall morphology. Proc. R. Soc. Lond. BBiol. Sci. 260, 85–89 (1995).

15. Nabity, P. D., Haus, M. J., Berenbaum, M. R. & DeLucia, E. H. Leaf-galling phylloxera on grapes reprograms host metabolism and morphology. PNAS 110, 16663–16668 (2013).

16. Soltis, P. S. et al. Floral variation and floral genetics in basal angiosperms. Am. J. Bot. 96, 110–128 (2009).

17. Mi, H. et al. PANTHER version 11: Expanded annotation data from gene ontology and reactome pathways, and data analysis tool enhancements. Nucleic Acids Res. 45, D183–D189 (2016).

18. Boss, P. K., Bastow, R. M., Mylne, J. S. & Dean, C. Multiple pathways in the decision to flower: Enabling, promoting, and resetting. Plant Cell 16, S18–S31 (2004).

19. Putterill, J., Laurie, R. & Macknight, R. It’Its time to flower: The genetic control of flowering time. Bioessays 26, 363–373 (2004).

20. Ausin, I., Alonso-Blanco, C. & Martinez-Zapater, J. M. Environmental regulation of flowering. Int. J. Dev. Biol. 49, 689–705 (2004).

21. Amasino, R. M. & Michaels, S. D. The timing of flowering. Plant Physiol. 154, 516–520 (2010)

22. Costantini, L., Battilana, J., Lamaj, F., Fanizza, G. & Grando, M. S. Berry and phenology-related traits in grapevine (*Vitis vinifera* L.): From quantitative trait loci to underlying genes. BMC Plant Biol. 8, 38 (2008).

23. Berardini, T. Z. et al. The Arabidopsis information resource: Making and mining the “gold standard” annotated reference plant genome. Genesis 53, 474–485 (2015).

24. UniProt Consortium. UniProt: The universal protein knowledgebase. Nucleic Acids Res. 45, D158–D169 (2016).

25. Bouché, F., Lobet, G., Tocquin, P. & Périlleux, C. FLOR-ID: An interactive database of flowering-time gene networks in *Arabidopsis thaliana*. Nucleic Acids Res. 44, D1167–D1171 (2015).

26. McClung, C. R., Lou, P., Hermand, V. & Kim, J. A. The importance of ambient temperature to growth and the induction of flowering. Front. Plant Sci. 7, 1266 (2016).

27. Boss, P. K. & Thomas, M. R. Association of dwarfism and floral induction with a grape ‘green revolution’ mutation. Nature 416, 847–850 (2002).

28. Blázquez, M. A., Soowal, L. N., Lee, I. & Weigel, D. LEAFY expression and flower initiation in Arabidopsis. Dev. 124, 3835–3844 (1997).

29. Yamaguchi, N., Jeong, C. W., Nole-Wilson, S., Krizek, B. A. & Wagner, D. AINTEGUMENTA and AINTEGUMENTA-LIKE6/PLETHORA3induce LEAFY expression in response to auxin to promote the onset of flower formation in Arabidopsis. Plant Physiol. 170, 283–293 (2016).

30. Van Dijk, A. D. & Molenaar, J. Floral pathway integrator gene expression mediates gradual transmission of environmental and endogenous cues to flowering time. Peer J. 5, e3197 (2017).

31. Hong, Y. & Jackson, S. Floral induction and flower formation - The role and potential applications of miRNAs. Plant Biotechnol. J. 13, 282–292 (2015).

32. Moon, J., Lee, H., Kim, M. & Lee, I. Analysis of flowering pathway integrators in Arabidopsis. Plant Cell Physiol. 46, 292–299 (2005).

33. Kardailsky, I. et al. Activation tagging of the floral inducer FT. Science 286, 1962–1965 (1999).

34. Kobayashi, Y., Kaya, H., Goto, K., Iwabuchi, M. & Araki, T. A pair of related genes with antagonistic roles in mediating flowering signals. Science 286, 1960–1962 (1999).

35. Samach, A. et al. Distinct roles of CONSTANS target genes in reproductive development of Arabidopsis. Science 288, 1613–1616 (2000).

36. Gocal, G. F. et al. GAMYB-like genes, flowering, and gibberellin signaling in Arabidopsis. Plant Physiol. 127, 1682–1693 (2001).

37. Achard, P., Herr, A., Baulcombe, D. C. & Harberd, N. P. Modulation of floral development by a gibberellin-regulated microRNA. Dev. 131, 3357–3365 (2004).

38. Ledger, S., Strayer, C., Ashton, F., Kay, S. A. & Putterill, J. Analysis of the function of two circadian-regulated CONSTANS-like genes. Plant J. 26, 15–22 (2001).

39. Weis, A., Walton, R., Crego C. L. Reactive plant tissue sites and the population biology of gall makers. Ann. Rev. Ent. 33, 467–486 (2003).

40. Carneiro, R. G., Isaias, R., Moreira, A. S. & Oliveira, D. C. Reacquisition of new meristematic sites determines the development of a new organ, the Cecidomyiidae gall on Copaifera langsdorffii Desf. (Fabaceae). Front. Plant Sci. 8, 1622 (2017).

41. Etchells, J. P., Provost, C. M., Mishra, L. & Turner, S. R. WOX4 and WOX14 act downstream of the *PXY* receptor kinase to regulate plant vascular proliferation independently of any role in vascular organisation. Dev. 140, 2224–2234 (2013).

42. Etchells, J. P. & Turner, S. R. The PXY-CLE41 receptor ligand pair defines a multifunctional pathway that controls the rate and orientation of vascular cell division. Dev. 137, 767–774 (2010).

43. Love, J. et al. Ethylene is an endogenous stimulator of cell division in the cambial meristem of Populus. PNAS 106, 5984–5989 (2009).

44. Jouannet, V., Brackmann, K. & Greb, T. (Pro)cambium formation and proliferation: Two sides of the same coin? Curr. Opin. Plant Biol. 23, 54–60 (2015).

45. Baima, S. et al. Negative feedback regulation of auxin signaling by ATHB8/ACL5-BUD2 transcription module. Mol. Plant 7, 1006–1025 (2014).

46. Berleth, T., Scarpella, E. & Prusinkiewicz, P. Towards the systems biology of auxin-transport-mediated patterning. Trends Plant Sci. 12, 151–159 (2007).

47. Furuta, K. M., Hellmann, E. & Helariutta, Y. Molecular control of cell specification and cell differentiation during procambial development. Annu. Rev. Plant Biol. 65, 607–638 (2014).

48. Cheng, Y., Dai, X. & Zhao, Y. Auxin biosynthesis by the YUCCA flavin monooxygenases controls the formation of floral organs and vascular tissues in Arabidopsis. Genes Dev. 20, 1790–1799 (2006).

49. Weigel, D. & Nilsson, O. A developmental switch sufficient for flower initiation in diverse plants. Nature 377, 495–500 (1995).

50. Carmona, M. J., Cubas, P. & Martinez-Zapater, J. M. VFL, the grapevine FLORICAULA/ LEAFY ortholog, is expressed in meristematic regions independently of their fate. Plant Physiol. 130, 68–77 (2002).

51. Krizek, B. A. Auxin regulation of Arabidopsis flower development involves members of the AINTEGUMENTA-LIKE/PLETHORA (AIL/PLT) family. J. Exp. Bot. 62, 3311–3319 (2011).

52. Winter, C. M. et al. LEAFY target genes reveal floral regulatory logic, cis motifs, and a link to biotic stimulus response. Dev. Cell 20, 430–443 (2011).

53. Yoon, E. K. et al. Auxin regulation of the microRNA390-dependent transacting small interfering RNA pathway in *Arabidopsis* lateral root development. Nucleic Acids Res. 38, 1382–1391 (2010).

54. Nemhauser, J. L., Feldman, L. J. & Zambryski, P. C. Auxin and *ETTIN* in *Arabidopsis* gynoecium morphogenesis. Dev. 127, 3877–3888 (2000).

55. Nagpal, P. et al. Auxin response factors *ARF6* and *ARF8*promote jasmonic acid production and flower maturation. Dev. 132, 4107–4118 (2005).

56. Hyun, Y., Richter, R. & Coupland, G. Competence to flower: Age-controlled sensitivity to environmental cues. Plant Physiol. 173, 36–46. (2017).

57. Joly, D. et al. Expression analysis of flowering genes from seedling-stage to vineyard life of grapevine cv. Riesling. Plant Sci. 166, 1427–1436 (2004).

58. Lu, F. et al. Arabidopsis REF6 is a histone H3 lysine 27 demethylase. Nat. Genet. 43, 715–719 (2011).

59. Dreni, L. & Kater, M. M. MADS reloaded: Evolution of the *AGAMOUS* subfamily genes. New Phytol. 201, 717–732 (2014).

60. Carmona, M. J., Calonje, M. & Martínez-Zapater, J. M. The *FT/TFL1* gene family in grapevine. Plant Mol. Biol. 63, 637–650 (2007).

61. Pastore, J. J. LATE MERISTEM IDENTITY2 acts together with LEAFY to activate APETALA1. Dev. 138, 3189–3198 (2011).

62. Prunet, N. et al. *REBELOTE, SQUINT*, and *ULTRAPETALA1* function redundantly in the temporal regulation of floral meristem termination in *Arabidopsis thaliana*. Plant Cell 20, 901–919 (2008).

63. Reyes-Olalde, J. I., Zuñiga-Mayo, V. M., Montes, R. A. C., Marsch-Martínez, N. & De Folter, S. Inside the gynoecium: At the carpel margin. Trends Plant Sci. 18, 644–655 (2013).

64. Ray, A. et al. *Arabidopsis* floral homeotic gene *BELL (BEL1)* controls ovule development through negative regulation of *AGAMOUS* gene (AG). PNAS 91, 5761–5765 (1994).

65. Sridhar, V. V., Surendrarao, A. & Liu, Z. *APETALA1* and *SEPALLATA3* interact with *SEUSS* to mediate transcription repression during flower development. Dev. 133, 3159–3166 (2006).

66. Das, P. et al. Floral stem cell termination involves the direct regulation of *AGAMOUS* by *PERIANTHIA*. Dev. 136, 1605–1611 (2009).

67. Rijpkema, A. S., Vandenbussche, M., Koes, R., Heijmans, K. & Gerats, T. Variations on a theme: Changes in the floral ABCs in angiosperms. Semin. Cell Dev. Biol. 21, 100–107 (2010).

68. Ferrándiz, C. et al. Carpel development. Adv. Bot. Res. 55, 1–73 (2010).

69. Liljegren, S. J. et al. *SHATTERPROOF*MADS-box genes control seed dispersal in *Arabidopsis*. Nature 404, 766–770 (2000).

70. Pinyopich, A. et al. Assessing the redundancy of MADS-box genes during carpel and ovule development. Nature 424, 85–88 (2003).

71. Honma, T. & Goto, K. Complexes of MADS-box proteins are sufficient to convert leaves into floral organs. Nature 409, 525–529 (2001).

72. Ditta, G., Pinyopich, A., Robles, P., Pelaz, S. & Yanofsky, M. F. The *SEP4* gene of *Arabidopsis thaliana* functions in floral organ and meristem identity. Curr. Biol. 14, 1935–1940 (2004).

73. Rijpkema, A. S., Zethof, J., Gerats, T. & Vandenbussche, M. The petunia *AGL6* gene has a *SEPALLATA-Uke* function in floral patterning. Plant J. 60, 1–9 (2009).

74. Xu, Y.-Y. Activation of the *WUS* gene induces ectopic initiation of floral meristems on mature stem surface in *Arabidopsis thaliana*. Plant Mol. Biol. 57, 773–784 (2005).

75. Pelaz, S., Tapia-Lopez, R., Alvarex-Buylla, E. & Yanofsky, M. Conversion of leaves into petals in *Arabidopsis*. Curr. Biol. 11, 182–184 (2001).

76. Blázquez, M. A. & Weigel, D. Integration of floral inductive signals in *Arabidopsis*. Nature 404, 889 (2000).

77. Yamaguchi, N. et al. Gibberellin acts positively then negatively to control onset of flower formation in *Arabidopsis*. Science 344, 638–641 (2014).

78. Calonje, M., Cubas, P., Martínez-Zapater, J. M. & Carmona, M. J. Floral meristem identity genes are expressed during tendril development in grapevine. Plant Physiol. 135, 1491–1501 (2004).

79. Dreni, L. & Zhang, D. Flower development: The evolutionary history and functions of the *AGL6*subfamily MADS-box genes. J. Exp. Bot. 67, 1625–1638 (2016).

80. Medina, C. et al. Characterization of microRNAs from *Arabidopsis* galls highlights a role for miR159 in the plant response to the root-knot nematode *Meloidogyne incognita*. New Phytol. 216, 882–896 (2017).

81. Rosen, H. R. The development of the *Phylloxera vasatrix* leaf gall. Science 43, 216–217 (1916).

82. Yamaguchi, Y. L. et al. Root-knot and cyst nematodes activate procambium-associated genes in *Arabidopsis* roots. Front. Plant Sci. 8, 1195 (2017).

83. Favaro, R. MADS-box protein complexes control carpel and ovule development in *Arabidopsis*. Plant Cell 15, 2603–2611 (2003).

84. Murai, K. Homeotic genes and the ABCDE model for floral organ formation in wheat. Plants 2, 379–395 (2013).

85. Li, J., Jia, D. & Chen, X. *HUA1*, a regulator of stamen and carpel identities in *Arabidopsis*, codes for a nuclear RNA binding protein. Plant Cell 13, 2269–2281 (2001).

86. Schonrogge, K., Harper, L. J. & Lichtenstein, C. P. The protein content of tissues in cynipid galls (Hymenoptera: Cynipidae): Similarities between cynipid galls and seeds. Plant Cell Environ. 23, 215–222 (2000).

87. Hartley, S. E. The chemical composition of plant galls: Are levels of nutrients and secondary compounds controlled by the gall-former? Oecol. 113, 492–501 (1998).

88. Von Aderkas, P., Rouault, G., Wagner, R., Rohr, R. & Roques, A. Seed parasitism redirects ovule development in Douglas fir. Proc. R. Soc. Lond. B Biol. Sci. 272, 1491–89.

89. Dilcher, D. Major innovations in angiosperm evolution in *Plants in Mesozoic time: Morphological innovations, phylogeny, ecosystems* (ed. Gee, C. T.) 97–116 (Indiana University Press, Bloomington, 2010).

90. Tooker, J. F. & Helms, A. M. Phytohormone dynamics associated with gall insects, and their potential role in the evolution of the gall-inducing habit. J. Chem. Ecol. 40, 742–753 (2014).

91. Yamaguchi, N. et al. A molecular framework for auxin-mediated initiation of flower primordia. Dev. Cell 24, 271–282 (2013).

92. Matsoukas, I. G. Interplay between sugar and hormone signaling pathways modulate floral signal transduction. Front Genet. 5, 218 (2014).

93. Voordeckers, K., Pougach, K., Verstrepen, K.J. How do regulatory networks evolve and expand throughout evolution? *Curr*. Opin. Biotech. 34, 280–288 (2015).

94. Díaz-Riquelme, J., Lijavetzky, D., Martínez-Zapater, J. M. & Carmona, M. Genome-wide analysis of MIKCC-type MADS box genes in grapevine. Plant Physiol. 149, 354–369 (2009).

95. Hawkins, C. & Liu, Z. A model for an early role of auxin in *Arabidopsis* gynoecium morphogenesis. Front. Plant Sci. 5, 327 (2014).

96. Sterling, C. Ontogeny of the phylloxera gall of grape leaf. Am JBot 39, 6–15 (1952).

97. Ghosh, S. & Chan, CK. Analysis of RNA-seq data using TopHat and Cufflinks. Methods Mol. Biol. 1374, 339–361 (2016).

